# Phylogenetic analysis shows canine distemper virus outbreak in stray dogs possibly occurs through spillover from wild carnivore reservoirs

**DOI:** 10.1101/2023.03.05.531168

**Authors:** Prajwol Manandhar, Rajindra Napit, Saman M Pradhan, Pragun G Rajbhandari, Jessie A Moravek, Pranav R Joshi, Rima D Shrestha, Dibesh Karmacharya

## Abstract

Canine distemper is a highly contagious, often fatal disease caused by canine distemper virus (CDV) in domestic dogs and wild carnivores. The virus has caused mass epidemics in wild carnivores of high conservation value such as tigers, lions and leopards in both wild and captivity. Hence, understanding and managing CDV outbreaks is particularly important in Nepal, which is home to many species of threatened wild carnivores including tigers, leopards, snow leopards, dholes and wolves, as well as a large population of stray dogs. Previous studies have suggested that CDV may pose a threat to wild carnivores, but there has not been any studies characterizing the genetic strains of the virus circulating in Nepal’s carnivores. We collected invasive and non-invasive biological samples from stray dogs in Kathmandu Valley and genetically characterized the strains of CDV in the dogs to belong to Asia-5 lineage by using phylogenetic analysis. The same lineage also contained CDV strains isolated from dogs, civets, red panda and lions in India. Based on our phylogenetic analysis, we think it is likely that in Nepal CDV is maintained through sylvatic cycle among small carnivore guilds allowing the recurring spillovers and outbreaks among free-ranging stray dogs and possibly large carnivores. It is crucial to prevent the virus transmission from reservoir hosts to other species, especially threatened populations of large carnivores in Nepal. Hence, we recommend for regular surveillance of CDV targeting small wild carnivores as well as vaccination programmes to control the disease spillover in stray dogs.

## Introduction

Canine distemper is a highly contagious, often fatal disease caused by canine distemper virus (CDV). CDV is a single-stranded enveloped RNA virus belonging to the *Morbillivirus* genus of Paramyxoviridae family [1]. The disease is transmitted by aerosol and causes characteristic respiratory, gastrointestinal and nervous symptoms in infected domestic dogs (*Canis lupus familiaris*) and wild carnivores [1-3]. While often considered to be a dog disease, CDV has been reported in almost all members of the Carnivora order, as well as in some primates and ungulates. Importantly, CDV has been known to cause mass epidemics in wild carnivores of high conservation value. For example, CDV outbreaks have caused mass mortality of African lions (*Panthera leo leo*) in Serengeti National Park and Asiatic lions (*Panthera leo persica*) in Gir National Park [4-6]. Similarly, CDV has been isolated from deceased Amur tigers (*Panthera tigris altaica*) and Amur leopards (*Panthera pardus orientalis*) in Eastern Russia [7, 8]. The disease has also been found to infect captive tigers and leopards in zoos in the United States, as well as farmed animals like minks, ferrets, martens in the United States and Europe [9, 10]. In recent years, these outbreaks have indicated that emerging lineages of CDV have expanded the host range of the disease.

Understanding and managing CDV outbreaks is particularly important in Nepal, which is home to many species of threatened wild carnivores including tigers, leopards, snow leopards, dholes and wolves, as well as a large population of stray dogs [11]. Previous studies have shown that CDV is prevalent in stray dog populations across Nepal. A study near Chitwan National Park in 2017 identified seroprevalence of CDV antibodies in 17% of stray dogs [12]. Similarly, a study in Manang in 2018 identified CDV antibodies in 70% of dogs, while 13% tested positive in RT-PCR screening [13]. In both situations, because the dogs had never been vaccinated for CDV, the presence of CDV antibodies indicated that these animals had been exposed and/or infected to the virus. Furthermore, Chitwan and Manang are both located in or near important Nepali wildlife regions, and the proximity of infected dogs and wild carnivore populations creates the potential for disease transmission.

Although these previous studies suggest that CDV may pose a threat to wild carnivores, no studies have genetically characterized local virus strains in Nepal. To better understand evolutionary origin of CDV in Nepal, we genetically characterized CDV from outbreaks in stray dogs in the Kathmandu Valley in 2017. We also inferred the possible ongoing sylvatic cycle of CDV circulation among carnivores in Nepal, based on phylogenetic analysis of CDV lineages across diverse hosts by analyzing publicly available sequences from domestic and wild CDV hosts around the world.

## Materials and Methods

### Study design and study site

We conducted this study to screen stray dogs for CDV in the Kathmandu Valley (Figure 1). We chose a densely populated locality in the Bhaktapur district that is adjoined by the Suryabinayak community forest on south (Figure 1). The Suryabinayak community forest is a contiguous forest patch in the southern outskirts of the Kathmandu Valley, and many wild carnivores like leopards, civets, martens and small felids can be found to occur in the forested hills [14]. We sampled free-ranging stray dogs in a dense settlement area in central Bhaktapur and used a stratified sampling scheme to collect a proportional number of samples from all parts of the region.

**Figure 1.**
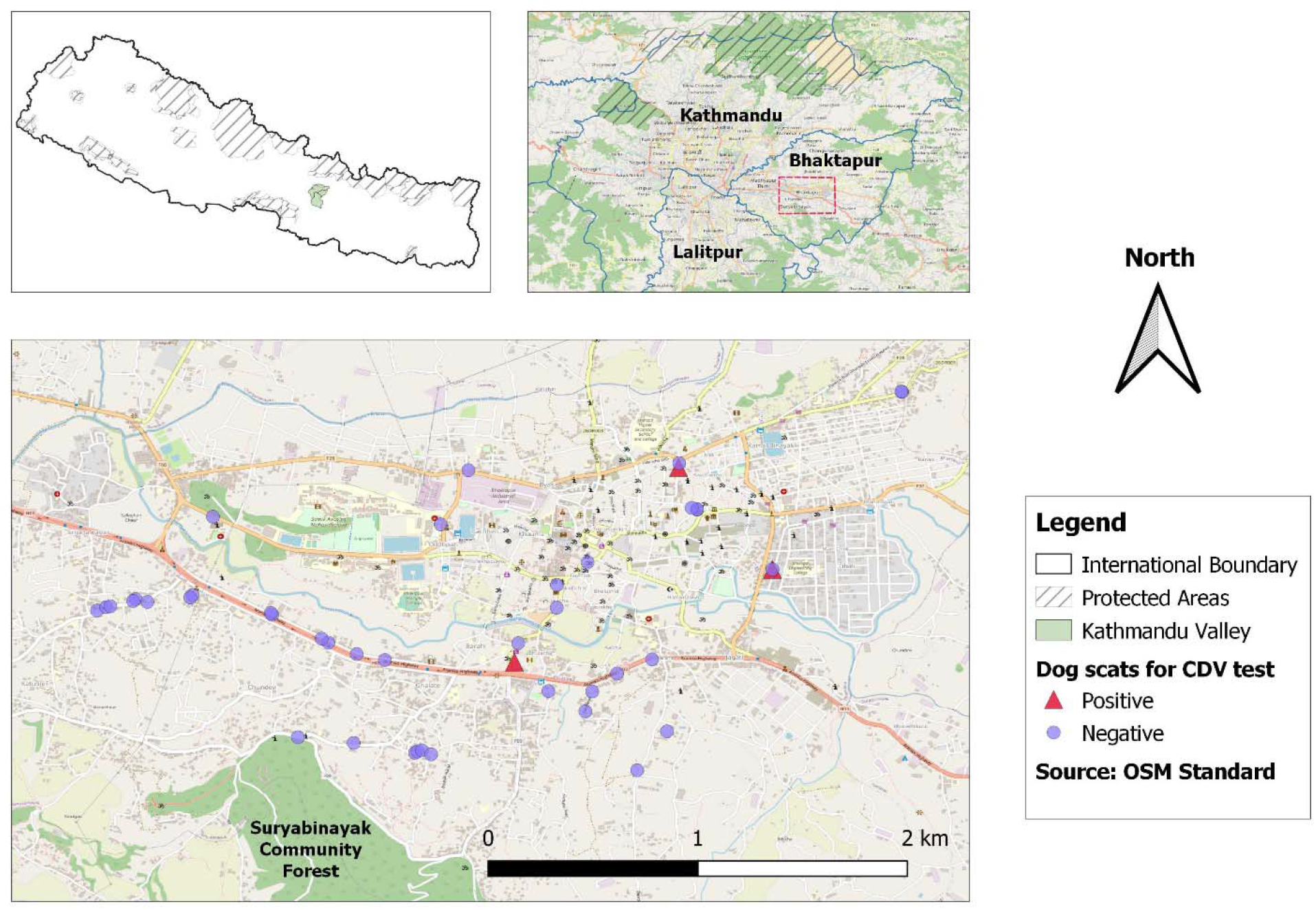
The locality in the center of Bhaktapur district, where dog scat samples were collected for detecting and characterizing CDV strains in stray dogs of Kathmandu Valley.

### Sample collection

We partnered with local veterinary clinics in Kathmandu and Bhanktapur to collect invasive biological samples from stray dogs in our sampling area. Local veterinary practitioners provided archived samples (ocular, saliva, fecal or rectal swabs) of stray dogs, suspected by veterinarians to have a CDV infection. The dogs either demonstrated clinical symptoms of CDV, including red eyes, cracked paws, nasal and ocular discharge, seizures or bodily twitching [eg. 15].

We also collected non-invasive fecal samples from stray dogs in central Bhaktapur (Figure 1). We sampled fresh dog scats in and along streets between 7 and 9 AM before the municipality cleaned the streets. The sample collector wore gloves and a facemask to prevent sample contamination and changed gloves between each sampling. We used a sterile cotton swab to dab the outer mucus layer of the scat and swirled in a cryovial filled with 0.5ml of Trizol stabilizer. We stored the samples in a -80°C freezer in Intrepid Nepal lab until further processing.

### Molecular screening of CDV by RT-PCR

We vortexed scat samples to homogenously distribute cells in the Trizol suspension. We performed RNA extraction using the Direct-zol RNA MiniPrep kit (Zymo Research, USA). We carried out cDNA synthesis from RNA elute using Invitrogen Superscript III (ThermoFisher, USA) and stored in -80°C freezer until downstream processing.

We performed PCR screening of CDV *phosphoprotein* gene on the cDNA using a *Morbillivirus* specific primer set that amplifies the ∼390 bp region of the gene [16]. We visualized PCR amplifications in 1.5% Agarose gel electrophoresis and sequenced positively amplified PCR products to confirm CDV.

### Phylogenetic analysis of CDV strains

After confirming presence of CDV in samples using the *phosphoprotein* gene, we targeted partial *hemagglutinin* gene sequence spanning ∼852 bp region in CDV positive samples for variant characterization. The *hemagglutinin* gene encodes for the viral surface glycoprotein that mediates fusion between the virus and the infected host cells. This gene is subjected to higher genetic variability and is therefore used as marker for lineage classification of CDV in most studies [17].

We utilized a tiled amplicon method to amplify and sequence two fragments (towards the 3’ region) using primers 3F/3R and 4F/4R of the *hemagglutinin* gene as provided in A Müller, E Silva, N Santos and G Thompson [18], each of which generated 563bp and 572bp respectively with about ∼40bp overlap regions. We sequenced each fragment from both directions and assembled fragments into a contiguous sequence (∼852bp) for further analysis.

To characterize the lineage of the CDV strains from stray dogs in Kathmandu, we compared the sample sequences collected for our study with all currently known lineages of CDV. We compiled a dataset of 245 full-length CDV *hemagglutinin* reference gene sequences (hereafter, referred as reference sequence) originating from at least 26 different countries across five continents (except Australia and Antarctica) collected from 1940 to 2018 (S1 Table). The compiled dataset included sequences from 17 different geographically defined lineages from host species belonging to the Canidae, Felidae, Mustelidae, Ailuridae, Procyonidae, and Ursidae families of carnivores, which are publicly available in the NCBI GenBank database. We aligned all the reference sequences along with sample sequences using MUSCLE ver3.8.425 [19], and visually inspected alignment and then trimmed and edited the sequences wherever required using AliView ver1.26 [20]. For the phylogenetic tree reconstruction, we selected the best-fit nucleotide substitution model (TPM1uf+G) based on Bayesian Information Criterion in jModeltest2 ver2.1.8 [21] from the alignment dataset. We conducted phylogenetic analysis using this model and the Bayesian inference method in MrBayes v3.2.7 [22] with 2,000,000 iterations, sampling every 2,000 iterations and discarding the first 25% as a burn-in. We visualized and annotated the phylogenetic tree using FigTree v1.4.4 [23].

## Results

### Molecular detection

We obtained clinical samples from eight symptomatic dogs admitted at two veterinary clinics in Bhaktapur and Kathmandu districts. From the eight dogs, we obtained a total of 15 CDV samples including ocular (n=7), rectal (n=7) and saliva (n=1) samples. Seven of the dogs tested positive for CDV via one or more of these sample types, while one individual dog tested negative in both ocular and rectal samples. Of the fifteen total samples taken, four ocular and four rectal samples tested positive. We also collected 44 scat samples from the streets of Bhaktapur, among which only three samples (6.8%) tested positive for CDV via *P-gene* PCR screening.

### Lineage characterization and hosts diversity of CDV

We determined *H-gene* partial sequences (∼800bp) from five CDV samples (four clinical and one scat source) isolated from dogs in the Kathmandu Valley (S2 Table). From the phylogenetic analysis, the CDV strains from dogs in Kathmandu Valley were classified as Asia-5 lineage. This lineage also contained CDV strains isolated from dogs, civet, red panda and Asiatic lions in India.

We found that CDV lineages were mainly clustered according to geographic regions. Each lineage consisted of strains isolated from diverse carnivore species co-occurring in the respective geographic regions. Virus strains isolated from carnivores belonging to Canidae and Mustelidae were the most commonly occurring ones across the lineages. The Asia-5 lineage where our Kathmandu Valley dog samples clustered is a sister clade to Africa-2 lineage that consisted of strains isolated from carnivores in the Serengeti landscape including African lions, African wild dogs and domestic dogs.

## Discussion

### CDV in the Kathmandu Valley

Our study genetically characterized CDV strains in dogs of Kathmandu as the Asia-5 lineage, which was first identified in 2016, described by M Bhatt, K Rajak, S Chakravarti, A Yadav, A Kumar, V Gupta, V Chander, K Mathesh, S Chandramohan and A Sharma [2] from dogs in India. Later, the same lineage was also found to be the causative variant behind fatal outbreaks in Asiatic lions in Gir National Park, Gujarat of western India during 2018, which caused the death of nearly two dozens of lions [6]. The same strains were also identified in Asian palm civet and red panda elsewhere in India [6, 24]. The origin of CDV strains found in Kathmandu dogs being the same as those found among carnivores in India, this indicates that the Asia-5 variant is prominent among reservoir hosts in Indian subcontinent region.

Similar regional patterns have been identified in other CDV strains in other regions. For example, the Africa-2 variant was first identified to be responsible for death of over 1,000 African lions in Serengeti National Park, Tanzania in 1985 [4]. The same strain was later isolated from feral dogs, spotted hyaena and bat-eared fox during investigation of the outbreak, and it was again detected in African wild dogs and jackals decades later [25]. Similarly, the Artic-like variant of CDV that was isolated from deceased Amur tigers and Amur leopards in Siberia were also found among martens across the region during a long-term CDV surveillance study of wild carnivores in Russia [26].

These studies indicate that sympatric carnivores are usually infected by regionally circulating similar or closely related strains, which suggests that CDV in stray dogs in Kathmandu might pose conservation challenges for large carnivores in Nepal’s wilderness areas. The hills and forests surrounding the Kathmandu Valley are habitat for wild carnivores and the stray dogs frequently interact with wild carnivores as prey or competitors. These interactions may drive pathogen spillover between domestic dogs and wild carnivore populations. Although wild carnivores have not been surveyed for CDV, spillover events from stray dog populations are highly possible and pose a potential threat to wild carnivores.

### Phylogenetic Analysis of CDV

As observed from our CDV phylogenetic tree (Figure 2), in most lineages domestic dogs are the most common CDV host or reservoir. Studies have suggested that CDV is mainly maintained among a group of wild carnivores, possibly small carnivore guilds such as martens, civets or mongooses that are relatively resilient in nature [26, 27], which later transmits to stray dogs from their interactions that occur when stray dogs attack or prey on the small carnivore species. This is also supported in our phylogenetic tree, where the second most common reservoir host among all lineages were mustelids. The species from Mustelidae family in our phylogenetic dataset consisted of European badger, Ferret, American mink, Yellow-throated marten, Beech marten, Fisher, European polecat, Sable, Siberian weasel and Asian badger (S1 Table). These mustelids are distributed mainly across Asia, Europe and North America. A comprehensive study by M Gilbert, N Sulikhan, O Uphyrkina, M Goncharuk, L Kerley, EH Castro, R Reeve, T Seimon, D McAloose and IV Seryodkin [26] suggested that mustelids in the Siberian wilderness may be maintaining the CDV at local level via sylvatic cycles across multi-host carnivore communities.

**Figure 2.**
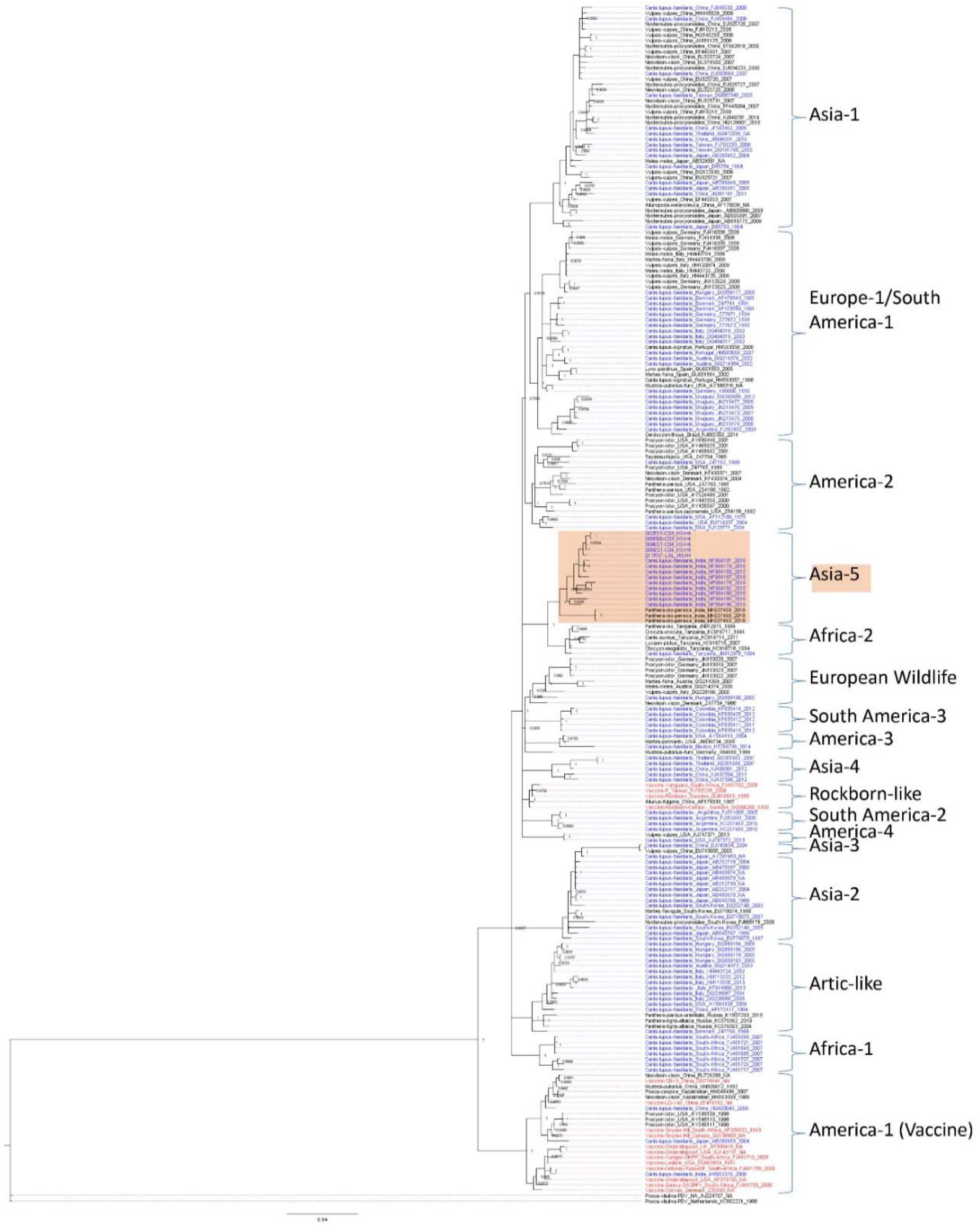
Phylogenetic tree constructed based on the 245 CDV hemagglutinin (H) gene isolates from 27 countries and 17 lineages, including the five samples collected from dogs (*Canis lupus familiaris)* in Nepal. Phylogenetic analysis was conducted using the Bayesian inference method in MrBayes with a run of 2,000,000 iterations, sub sampling per 2,000 iterations and discarding 25% of sample for burn-in. Common CDV vaccines were highlighted in red and two sequences of Phocine distemper virus were selected as the outgroup. Posterior probability of nodes representing significant or lineage defining branching points are indicated on the tree.

### Conservation Implications

In countries with large stray dog populations like Nepal, dogs have the potential to become CDV vectors as they frequently interact (attack/kill) with small carnivores like mustelids, and then fall prey to large cats like snow leopards, leopards and tigers. These ecological dynamics increase the chances of CDV spillovers and infection from small carnivores to stray dogs to large carnivore species, which are already threatened by a variety of anthropogenic factors [28]. The Kathmandu Valley is one such example where CDV spillover dynamics may be actively ongoing as rapid urbanization and anthropogenic encroachment of wildlife habitats are closing the gaps between forests and urban spaces. Studies in other regions of Nepal like Manang and Chitwan that also found a high prevalence of CDV antibodies in dogs suggest that this cycle may already be an ongoing process in most regions across the country. As such, more surveillances using molecular sequencing are required to confirm and characterize the CDV among suspected sick dogs and wild carnivores in Nepal. In addition to the carnivores, species such as ungulates and monkeys have been infected with CDV in the past and should also be surveilled [29].

Based on our phylogenetic analysis, we think it is likely that in Nepal CDV is maintained through sylvatic cycle among small carnivore guilds allowing the recurring spillovers and outbreaks among free-ranging stray dogs and possibly large carnivores. We stress on Nepal’s CDV surveillance programmes to expand from stray dogs to small carnivores such as martens, badgers, weasels and otters, as well as large carnivores. This study suggests that the unmanaged and growing populations of stray dogs in urban and rural areas are important drivers of the disease and pathogen transmission and pose a threat to wildlife. Thus, specific policies and action plans are utmost required to be implemented in order to prevent such outbreaks as well as reduce the interactions between dogs and wild carnivores in Nepal.

## Declarations

### Ethics approval and consent to participate

The authors affirm that the Journal’s ethical standards, as stated on the author guidelines page, were upheld throughout the study. Samples of dogs were obtained from private veterinary clinics in the Kathmandu and Bhaktapur districts of Nepal that had brought the dogs in for treatment of suspected illnesses, and no dogs were specifically procured for the study. Permission to collect dog scat samples in Bhaktapur was granted by the relevant government departments and municipality.

## Consent for publication

Not applicable.

## Availability of data and materials

All the data are included within the manuscript. The nucleotide sequences generated in this study are deposited in NCBI Genbank and accessions listed in Supplementary Table S2.

## Authors’ contributions

PM, RN, JAM and DK designed the study. PM, RN, JAM, PRJ and RDS collected samples. RN and SMP performed lab analysis. PM and PGR performed bioinformatics analysis. PM, RN, PGR, JAM and RDS prepared and reviewed initial drafts. All authors critically reviewed and finalized the manuscript.

## Acknowledgements

We thank Department of Livestock Services, Department of National Park and Wildlife Conservation and Bhaktapur Municipality for providing us permission to conduct this study. We thank Dr Martin Gilbert for guiding us during lab assay development of CDV RT-PCR detection and sequencing. We express our gratitude to all staffs, interns, collaborators at CMDN, Intrepid Nepal and Vet for Your Pet Animal Hospital who supported during various phases in the project.

## References

1. Zhao J, Shi N, Sun Y, Martella V, Nikolin V, Zhu C, Zhang H, Hu B, Bai X, Yan X: Pathogenesis of canine distemper virus in experimentally infected raccoon dogs, foxes, and minks. Antiviral research 2015, 122:1–11.

2. Bhatt M, Rajak K, Chakravarti S, Yadav A, Kumar A, Gupta V, Chander V, Mathesh K, Chandramohan S, Sharma A: Phylogenetic analysis of haemagglutinin gene deciphering a new genetically distinct lineage of canine distemper virus circulating among domestic dogs in India. Transboundary and emerging diseases 2019, 66(3):1252–1267.

3. Krakowka S, Axthelm M, Johnson G, Olsen R, Krakowka S, Blakeslee J: Comparative pathobiology of viral diseases. In.: CRC Press, Boca Raton; 1985.

4. Roelke-Parker ME, Munson L, Packer C, Kock R, Cleaveland S, Carpenter M, O’Brien SJ, Pospischil A, Hofmann-Lehmann R, Lutz H: A canine distemper virus epidemic in Serengeti lions (Panthera leo). Nature 1996, 379(6564):441–445.

5. Cleaveland S, Appel M, Chalmers W, Chillingworth C, Kaare M, Dye C: Serological and demographic evidence for domestic dogs as a source of canine distemper virus infection for Serengeti wildlife. Veterinary microbiology 2000, 72(3-4):217–227.

6. Mourya DT, Yadav PD, Mohandas S, Kadiwar R, Vala M, Saxena AK, Shete-Aich A, Gupta N, Purushothama P, Sahay RR: Canine distemper virus in Asiatic lions of Gujarat State, India. Emerging infectious diseases 2019, 25(11):2128.

7. Quigley KS, Evermann JF, Leathers CW, Armstrong DL, Goodrich J, Duncan NM, Miquelle DG: Morbillivirus infection in a wild Siberian tiger in the Russian Far East. Journal of wildlife diseases 2010, 46(4):1252–1256.

8. Sulikhan NS, Gilbert M, Blidchenko EY, Naidenko SV, Ivanchuk GV, Gorpenchenko TY, Alshinetskiy MV, Shevtsova EI, Goodrich JM, Lewis JC: Canine distemper virus in a wild far eastern leopard (Panthera pardus orientalis). Journal of wildlife diseases 2018, 54(1):170–174.

9. Appel MJ, Yates RA, Foley GL, Bernstein JJ, Santinelli S, Spelman LH, Miller LD, Arp LH, Anderson M, Barr M: Canine distemper epizootic in lions, tigers, and leopards in North America. Journal of Veterinary Diagnostic Investigation 1994, 6(3):277–288.

10. Harder TC, Kenter M, Vos H, Siebelink K, Huisman W, Van Amerongen G, Örvell C, Barrett T, Appel M, Osterhaus A: Canine distemper virus from diseased large felids: biological properties and phylogenetic relationships. Journal of General Virology 1996, 77(3):397–405.

11. Adhikari RB, Shrestha M, Puri G, Regmi GR, Ghimire TR: Canine Distemper Virus (CDV): an emerging threat to Nepal’s wildlife. Applied Science and Technology Annals 2020, 1(1):149–154.

12. Sadaula A, Joshi JD, Lamichhane BR, Gairhe KP, Subedi N, Pokheral CP, Thapaliya S, Pandey G, Rijal KR, Pandey P: Seroprevalence of Canine Distemper and Canine Parvovirus Among Domestic Dogs in Buffer Zone of Chitwan National Park, Nepal. 2022.

13. Ng D, Carver S, Gotame M, Karmasharya D, Karmacharya D, Man Pradhan S, Narsingh Rana A, Johnson CN: Canine distemper in Nepal’s Annapurna Conservation Area– Implications of dog husbandry and human behaviour for wildlife disease. PloS one 2019, 14(12):e0220874.

14. Katuwal HB, Basent H, Sharma HP, Koirala S, Khanal B, Neupane KR, Thapa KB, Panta DB, Parajuli K, Lamichhane S: Wildlife assessment of the Chandragiri hills, Kathmandu: Potentiality for ecotourism. European Journal of Ecology 2020, 6(1):27–50.

15. Creevy KE: Overview of Canine Distemper - Generalized Conditions - Merck Veterinary Manual. In. Edited by Corp. MSD; 2016.

16. Barrett T, Visser I, Mamaev L, Goatley L, Van Bressem M-F, Osterhaus A: Dolphin and porpoise morbilliviruses are genetically distinct from phocine distemper virus. Virology 1993, 193(2):1010–1012.

17. Bolt G, Jensen TD, Gottschalck E, Arctander P, Appel MJ, Buckland R, Blixenkrone M: Genetic diversity of the attachment (H) protein gene of current field isolates of canine distemper virus. Journal of General Virology 1997, 78(2):367–372.

18. Müller A, Silva E, Santos N, Thompson G: Domestic dog origin of canine distemper virus in free-ranging wolves in Portugal as revealed by hemagglutinin gene characterization. Journal of wildlife diseases 2011, 47(3):725–729.

19. Edgar RC: MUSCLE: a multiple sequence alignment method with reduced time and space complexity. BMC bioinformatics 2004, 5(1):1–19.

20. Larsson A: AliView: a fast and lightweight alignment viewer and editor for large datasets. Bioinformatics 2014, 30(22):3276–3278.

21. Darriba D, Taboada GL, Doallo R, Posada D: jModelTest 2: more models, new heuristics and parallel computing. Nature methods 2012, 9(8):772–772.

22. Ronquist F, Teslenko M, Van Der Mark P, Ayres DL, Darling A, Höhna S, Larget B, Liu L, Suchard MA, Huelsenbeck JP: MrBayes 3.2: efficient Bayesian phylogenetic inference and model choice across a large model space. Systematic biology 2012, 61(3):539–542.

23. Rambaut A: FigTree v1. 4. In.; 2012.

24. Kodi H, Putty K, Ganji VK, Bhagyalakshmi B, Reddy YN, Satish K, Prakash MG: H gene-based molecular characterization of field isolates of canine distemper virus from cases of canine gastroenteritis. Indian Journal of Animal Research 2021, 55(5):561–567.

25. Nikolin VM, Olarte□Castillo XA, Osterrieder N, Hofer H, Dubovi E, Mazzoni CJ, Brunner E, Goller KV, Fyumagwa RD, Moehlman PD: Canine distemper virus in the Serengeti ecosystem: molecular adaptation to different carnivore species. Molecular ecology 2017, 26(7):2111–2130.

26. Gilbert M, Sulikhan N, Uphyrkina O, Goncharuk M, Kerley L, Castro EH, Reeve R, Seimon T, McAloose D, Seryodkin IV: Distemper, extinction, and vaccination of the Amur tiger. Proceedings of the National Academy of Sciences 2020, 117(50):31954–31962.

27. Kapil S, Yeary TJ: Canine distemper spillover in domestic dogs from urban wildlife. Veterinary Clinics: Small Animal Practice 2011, 41(6):1069–1086.

28. Adhikari B, Baral K, Bhandari S, Szydlowski M, Kunwar RM, Panthi S, Neupane B, Koirala RK: Potential risk zone for anthropogenic mortality of carnivores in Gandaki Province, Nepal. Ecology and Evolution 2022, 12(1):e8491.

29. Beineke A, Baumgärtner W, Wohlsein P: Cross-species transmission of canine distemper virus—an update. One Health 2015, 1:49–59.

