## Supplementary material for "Phylogenetic analysis shows canine distemper virus outbreak in stray dogs possibly occurs through spillover from wild carnivore reservoirs": Table S2

S2 Table. List of hemagglutinin gene sequences of CDV generated in this study along with their details and Genbank accession numbers.

| **S.N.** | **Isolate ID** | **Isolation host** | **Isolation_source** | **Country** | **District** | **Collection date** | **Genbank accession** |
| --- | --- | --- | --- | --- | --- | --- | --- |
| 1 | D02EST-CD3 | *Canis lupus familiaris* | occular/rectal | Nepal | Bhaktapur | 2018-01-20 | OQ363404 |
| 2 | D04EST-CD4 | *Canis lupus familiaris* | occular/rectal | Nepal | Bhaktapur | 2018-01-20 | OQ363406 |
| 3 | D06EST-CD7 | *Canis lupus familiaris* | occular | Nepal | Bhaktapur | 2018-01-20 | OQ363407 |
| 4 | D09FED-CD3 | *Canis lupus familiaris* | feces | Nepal | Bhaktapur | 2018-03-27 | OQ363405 |
| 5 | D17RST-LAL | *Canis lupus familiaris* | ocular/rectal/saliva | Nepal | Kathmandu | 2018-04-03 | OQ363403 |
